# Predicting species distributions in the open ocean with convolutional neural networks

**DOI:** 10.1101/2023.08.11.551418

**Authors:** Gaétan Morand, Alexis Joly, Tristan Rouyer, Titouan Lorieul, Julien Barde

**Affiliations:** UMR Marbec, IRD, Univ Montpellier, CNRS, Ifremer; LIRMM, INRIA, Univ. Montpellier

**Keywords:** deep learning, megafauna, open ocean, pelagic species, species distribution models

## Abstract

As biodiversity plummets due to anthropogenic disturbances, the conservation of oceanic species is made harder by limited knowledge of their distributions and migrations. Indeed, tracking species distributions in the open ocean is particularly challenging due to the scarcity of observations and the complex and variable nature of the ocean system. In this study, we propose a new method that leverages deep learning, specifically convolutional neural networks (CNNs), to capture spatial features of environmental variables. This novelty eliminates the need to predefine these features before modelling and creates opportunities to discover unexpected correlations. Our aim is to present the results of the first trial of this method in the open ocean, discuss limitations and provide feedback for future improvements or adjustments.

In this case study, we considered 38 taxa comprising pelagic fishes, elasmobranchs, marine mammals, marine turtles and birds. We trained a model to predict probabilities from the environmental conditions at any specific point in space and time, using species occurrence data from the Global Biodiversity Information Facility (GBIF) and environmental data from various sources. These variables included sea surface temperature, chlorophyll concentration, salinity and fifteen others.

During the testing phase, the model was applied to environmental data at locations where species occurrences were recorded. The classifier accurately predicted the observed taxon as the most likely taxon in 69% of cases and included the observed taxon among the top three most likely predictions in 89% of cases. These findings show the adequacy of deep learning for species distribution modelling in the open ocean.

Additionally, this purely correlative model was then analysed with explicability tools to understand which variables had an influence on the model’s predictions. While variable importance was species-dependent, we identified finite-size Lyapunov exponents (FSLEs), sea surface temperature, pH and salinity as the most influential variables, in that order. These insights can prove valuable for future species-specific ecology studies.

## 1. Introduction

### 1.1 Background

The open ocean is a vast and complex ecosystem that covers over 70% of the Earth’s surface, yet it remains one of the least understood and studied ecosystems on our planet [1, 2]. It plays a critical role in regulating the Earth’s climate and biogeochemical cycles (including nutrient cycles and carbon sequestration), making it a vital component of all life on Earth [3, 4].

However, the ocean is facing a range of human-induced threats, including over-fishing, pollution and climate change [5, 6, 7]. These threats can have serious consequences for marine biodiversity and therefore negatively impact the livelihoods of millions of people who depend on the oceans for their food or income [8].

To solve these most pressing challenges, a necessary first step is to understand how marine life is distributed within the open ocean. Species distribution models can provide valuable insights into where different species are likely to be found and how environmental factors drive their distribution [9]. By developing accurate and reliable models, we can identify areas that are most threatened by foreseen local disturbances and develop effective conservation strategies to protect these ecosystems.

Furthermore, changes in the Earth’s climate are already affecting ocean conditions, namely warming waters, ocean acidification and sea level rise, among others [10]. This makes it even more urgent to understand the link between environmental variables and species distributions, to be able to predict how marine biodiversity may respond to these changes. This information is critical for informing decision-making and management efforts to ensure the long-term sustainability of marine ecosystems and the services they provide to society.

Therefore, studying species distribution in the open ocean is essential for advancing our understanding of these complex ecosystems and for developing effective conservation and management strategies to protect them.

### 1.2 Existing methods for predicting species distributions

A wide variety of Species Distribution Models (SDMs) have been discussed in literature [11]. This is generally done through modelling a species-specific *environmental niche* where environmental conditions are favourable to the species in the long term, shaped by natural selection [12]. Predictors are chosen empirically to try and predict the species’ ranges from observed species’ occurrences.

Usually, SDMs use climatological summaries of environmental data, at the location where the observation is recorded. These spatially isolated data are unable to convey the full nature of the environmental seascape around animals, as single values cannot represent more complex bathymetry features such as trenches for example. The same applies to other variables, which spatial structure may be more important than isolated values: algal blooms, temperature fronts, eddies, etc. Yet these spatial structures represent processes which are essential to ascertain species distributions [13, 14].

A solution to this shortcoming is to include the environmental data in the neighbourhood of species occurrences, but the number of predictors then becomes much larger than the number of observations. This is unfit for statistical models and requires a feature extraction step to summarize input data into fewer significant variables. This work may be carried out manually, which enables the model to take advantage of scientists’ expert knowledge. This is how some spatial features are added into SDMs [15], but it limits the performance of the model to the scope of existing knowledge and prevents the discovery of previously unknown influential factors.

Furthermore, the use of climatological summaries prevents taking advantage of these spatial features, as they are lost when averaging the values over time. While some climatology products try to mitigate this shortcoming (*e.g.* frequency of presence of fronts [16]), countless such products would be necessary to capture all types of features. Therefore, the only remaining solution is to use instantaneous values at the time of species occurrence [17]. This slightly changes the objective of the model: it does not try to model the ecological niche anymore, but dynamic distributions of the species and becomes a dynamic SDM [18]. This development is necessary if we want to include the aforementioned spatial structures into predictors. Incidentally, this makes the predictions highly dependent on time, which has two unintended benefits: **1.** making it possible to include the variability of environmental data into SDM predictors, which has been identified as a way to improve their performance [19] and **2.** allowing modelling the variations of distributions over time, which is especially interesting for highly mobile species or those that have rapid population dynamics [20, 21].

This calls for new methods to extend the scope of SDMs to fully take into account the complex spatial structures of environmental seascapes and, as a direct benefit, their variability over time.

### 1.3 Potential benefits of using deep learning for modelling marine species distribution

Convolutional neural networks (CNNs) were designed for image processing, so they have embedded feature extractors that are designed to detect multiple levels of details using convolution layers [22]. As the training advances deeper within the layers, small details are increasingly pooled together to be able to detect much more complex shapes. With image classification, one can identify the following levels, from most precise to coarser:

1. Values of specific pixels
2. Value of a small group of pixels: textures, edges
3. Association of several groups of pixels: shapes, geometric features
4. Association of several shapes: objects, animals, plants
5. Average and extreme values on the whole image: brightness/tint

This is especially useful with environmental data raster layers (from satellite observations or models) as it enables the model to detect the same various levels of details on environmental variables. Here are some examples of the same levels of detail, applied to environmental variables:

1. Values at a given point
2. Homogeneity of the variable in the neighbourhood of occurrence: fronts, slopes
3. Geographic features: bays, underwater canyons, river plumes
4. Complex shapes: current structures, cyclones
5. Average and extreme values over the buffer zones

The use of CNNs to model species distributions was successfully developed for terrestrial plants [23]. The CNN architecture proved especially useful to capture spatial features, as well as to transfer knowledge from better-known species to lesser-known species [24]. CNN-based SDMs, as described here, usually predict species distributions by providing a classification rather than regressions.

While these studies were mostly based on satellite imagery (Sentinel-2), optical data is not enough to represent the state of the oceanic environment, although it yields some interesting products such as chlorophyll concentration. These products and many other significant oceanic variables are available as processed data sets, which should be used for a more comprehensive view of oceanic conditions. Another difference with this previous work is the high temporal variability of the oceanic seascape. Here we present an adaptation of the work of Botella et al. [23] and Deneu et al. [24] that includes these adaptations to the specificity of the open ocean.

### 1.4 Objectives of the study

Through the present study, we explore the possibilities that deep-learning-based SDMs offer in the open ocean. We first give a detailed overview of the data that was used to build our model. Then we show the results that we obtained, including performance metrics and distribution maps. Finally, we point out the limits that we have found with our methodology choices and suggest ways to improve the results’ quality in the future.

## 2. Methods

The main step of our process is to build a model to relate species presence to environmental data. To achieve this, we used occurrence data from the Global Biodiversity Information Facility (GBIF) [25] and downloaded environmental data in a buffer around each of their locations, at the date of their occurrence. All the data sets are freely available.

It is important to note that as training data is presence-only, we cannot predict abundance or any absolute measure of presence. That’s why we modelled a multivariate output, where predictions are observation probabilities, relative to the 38 taxa that are the subject of this study. After training, this provided us with a model which takes environmental data as input and outputs a vector of observation probabilities (one for each taxon). The full process is summarized in Figure 1 and is explained in detail in this section.

**Figure 1.**
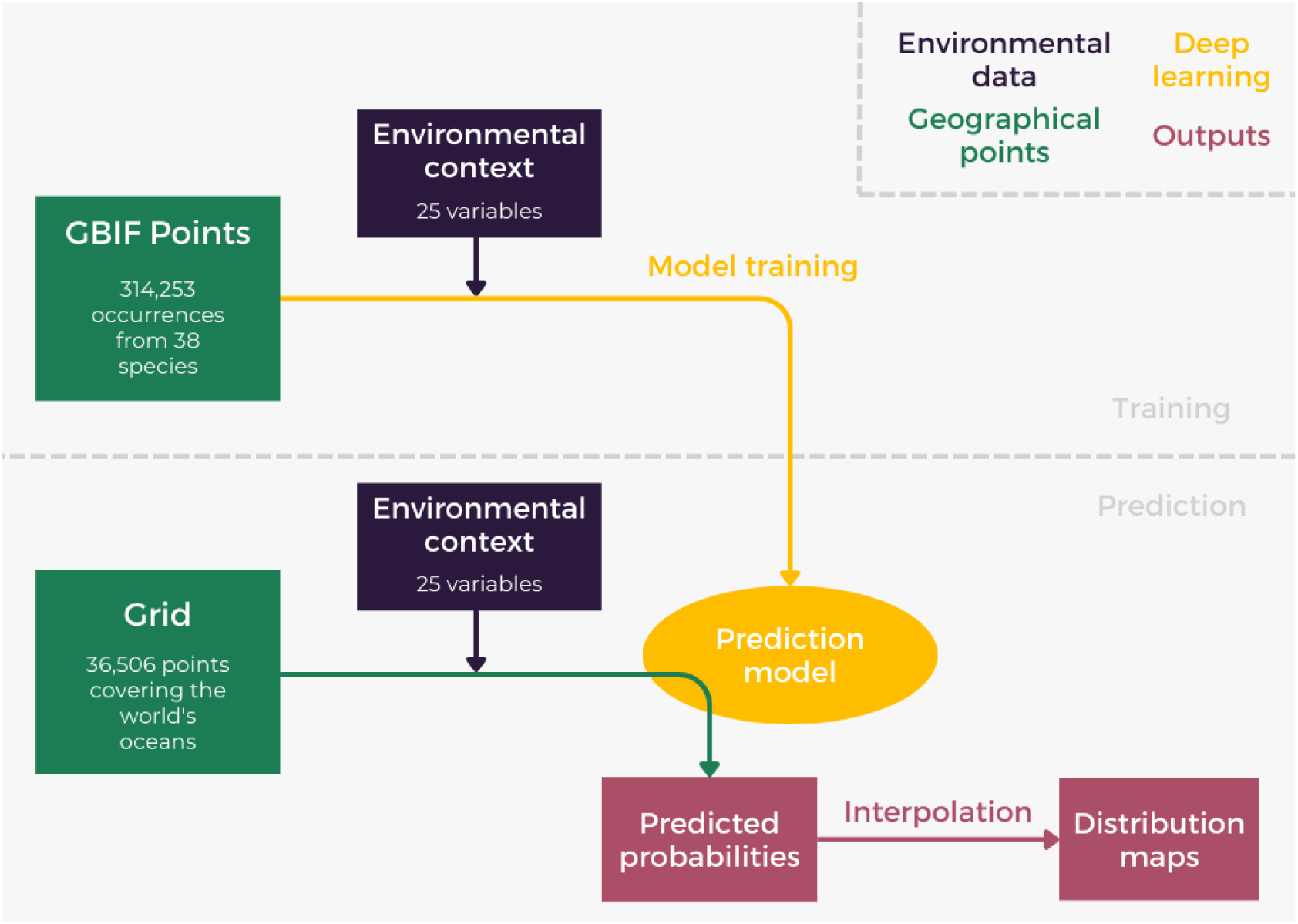
Summary view of the analysis process: model training in the top half and predictions in the bottom half.

### 2.1 Description of the occurrence data

Thirty-eight marine species or genera were selected for the proof of concept that is described in the present article. They include large pelagic fishes, elasmobranchs, turtles, sea mammals and two species of marine birds (see Table 1). They were chosen based on the availability of occurrence data, and special attention was given to their diversity in order to stress-test our model. The sample contains highly mobile and sessile species, widespread and local ones, some which live in biodiversity hotspots and others in less frequented waters. While sessile species (*Acropora*) cannot move in response to changing environmental conditions, the model may learn suitable conditions from geographical or long-term patterns, which could be useful to study the potential impact of temporary episodes (*e.g.* El Niño Southern Oscillation) or slower trends (*e.g.* ocean warming).

**Table 1.**
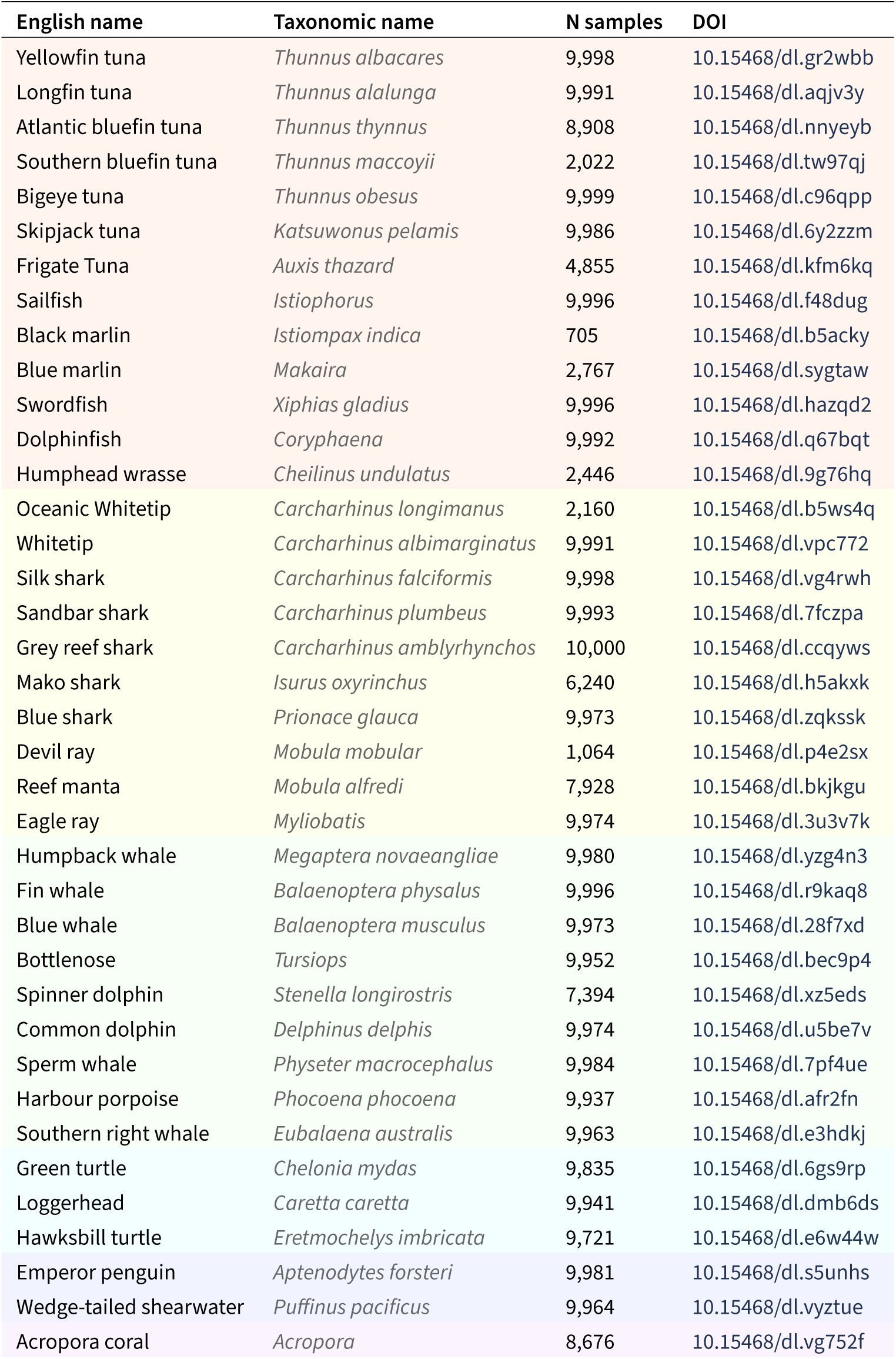
Species that were included in the study, coloured by taxonomic class. The last column is the digital object identifier (DOI) of downloaded archives.

Presence data for these taxa were downloaded from the GBIF. This database contains species occurrences that are free to access and download, which is essential for reproducibility. Some flawed data is unavoidably present in the database, but small errors in the geographical coordinates are not a problem as the oceanographic landscape that we consider has limited precision due to environmental data resolution (see Table 2). Furthermore, convolutional neural networks are known to be robust against occasional labelling mistakes [26].

**Table 2.**
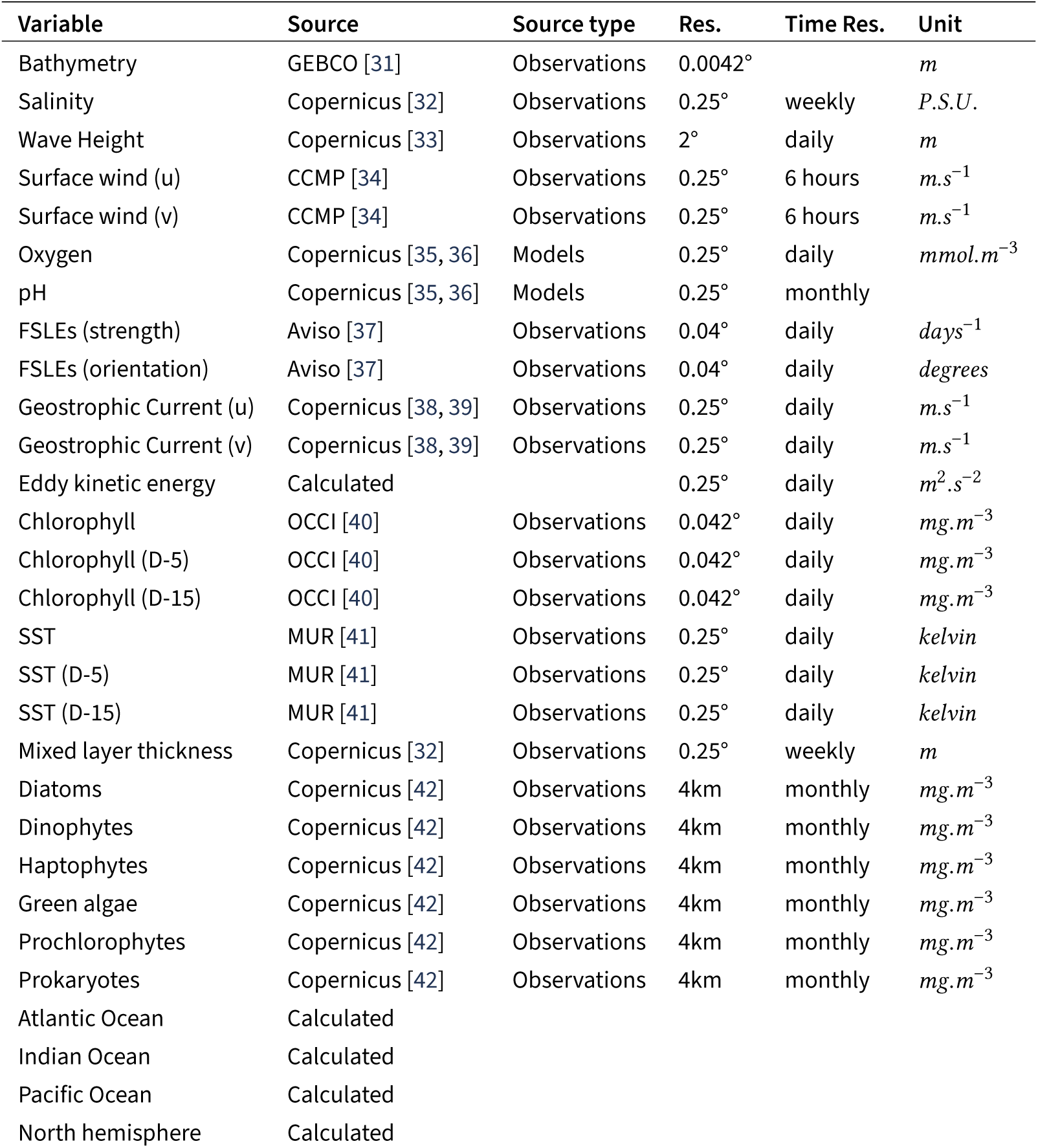
29 layers used as input data (re-ordered for clarity). NB: Eddy kinetic energy is calculated using geostrophic current data. Res. = Resolution. P.S.U. = Practical salinity Unit.

Digital object identifiers (DOIs) for the download of each species are available in Table 1. When more than 10,000 occurrences of a taxon were available, a random sample of 10,000 occurrences was selected. In all cases, GBIF identifiers of occurrences that were actually used are available in the training data set CSV files (*id* column). We removed points located on the continents and no other filtering was conducted on geographical precision.

We made the debatable choice not to intervene in the input data, as we cannot assume any consistent rules over all data sets. For example duplicates might be multiple sightings or simply mistakes. This choice is arbitrary and more thorough data cleaning could be beneficial to future work.

This added up to 314,253 occurrences for all taxa, depicted in Figure 2.

**Figure 2.**
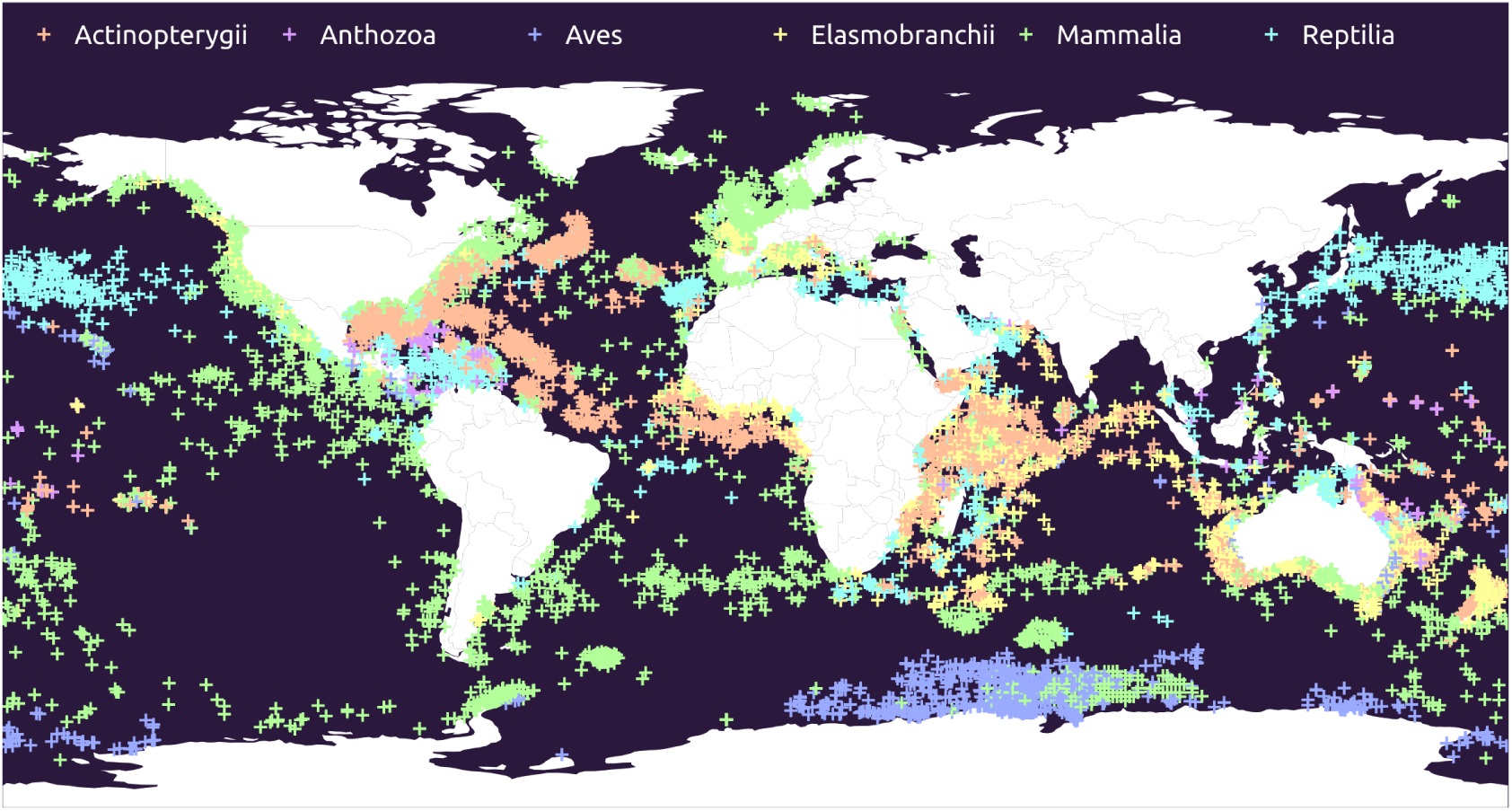
Random sample (10%) of the training data set, coloured by taxonomic class.

### 2.2 Description of the environmental data used as inputs

Eighteen environmental variables were considered, some from satellite observations and others from models. Three of them contain two values: both strength and orientation components (*i.e.*, polar coordinates) for finite-size Lyapunov exponents (FSLEs), and both zonal and meridional components for surface wind and geostrophic current. Temperature and chlorophyll values were also included 15 and 5 days before the occurrences, as it was previously demonstrated that marine animals may have a delayed response to some variables, especially temperature [27]. Finally, four geographical variables were added (see Section 2.3.2). Even though these four variables were constant over each patch, they were encoded as layers with equal dimensions to the other variables. This was done for simplicity of implementation, as well as to take advantage of GPUs’ efficiency at parallelized computation. Overall, this amounts to a total of 29 layers of data, shown in Table 2.

Each of these layers is two-dimensional: for 3D data only the surface layer was downloaded. However all the tools that we use in this study are compatible with 3D environmental data. We chose to focus on other aspects in this study, but the vertical dimension may be included into future work.

### 2.3 Data preparation

#### 2.3.1 Enrichment

Environmental data were downloaded in a buffer around the occurrences using the GeoEnrich python package, which was developed for this purpose and is made available to other researchers for a wide range of uses in the GitHub IRDG2OI/geoenrich repository [28]. The package implements caching, so that it does not make requests to the server when data has been downloaded already.

Data was downloaded for the closest available date to the occurrence. A spatial buffer of 115 km was used, to include at least one data point from the least precise data (2° resolution). This is consistent with values of daily potential movement for fast animals that may travel up to 120 km per day [29, 30]. All the available data within this buffer were downloaded into arrays.

These data arrays with various resolutions (minimum 1 × 1 for wave height, maximum 493 × 493 for bathymetry) also have various horizontal dimensions due to the longitude contraction closer to the poles. They were all interpolated (up-scaled or down-scaled depending on the initial resolution) to fit the same 32 × 32 grid centred around the occurrence. This grid has a resolution of approximately 7 km.

#### 2.3.2 Ocean basin and hemisphere

An initial goal of the study was to produce a geography-agnostic model, which means that two points with the same oceanic conditions, wherever they are, should yield the same predictions. But this is ecologically wrong for one main reason: natural barriers prevent animals from navigating anywhere in the long term, namely continents and for some species, the warm waters around the equator.

Because we propose a type of dynamic SDM, we cannot capture these long-term barriers, so we have to include them artificially. Therefore we added four binary variables: three for the main oceanic basins and one for the hemisphere.

The world’s oceans were split into three main basins: the Atlantic, Indian and Pacific oceans. Very few of the occurrences were located in the Arctic Ocean: they were assigned the closest of these ocean basins. Occurrences from the Southern Ocean were more numerous and are not separated from these three oceans by any physical barrier, so they could be assigned to the closest one.

It is important to note that the Ocean basin and the hemisphere are the only geographical information provided to the model. This is by design to avoid learning the observation bias that is present in the training data.

#### 2.3.3 Feature scaling

All data were scaled to the [0, 1] interval and saved into a data cube (32 × 32 geographical pixels × 29 layers). Outliers (highest and lowest 1% of the values of the training data set) were replaced with the corresponding extrema and the scaling factors were saved to be reapplied to any subsequent input data.

Some data are missing because of natural phenomena such as clouds, or because the occurrences were out of the data set time range. In that case, we used the median value of the variable over the tile. If data was missing over the whole tile, we used the median value over the whole data set instead. This does not allow the model to differentiate unobserved data (*e.g.* because of clouds) from no-data areas (*e.g.* coast), but this is not an issue as land pixels are already explicitly provided in the bathymetry layer.

Figure 3 shows an example of all the data that are included in the data cube used for training, with the feature scaling reversed in order to show the real values. The figure does not show the four binary geographical variables.

**Figure 3.**
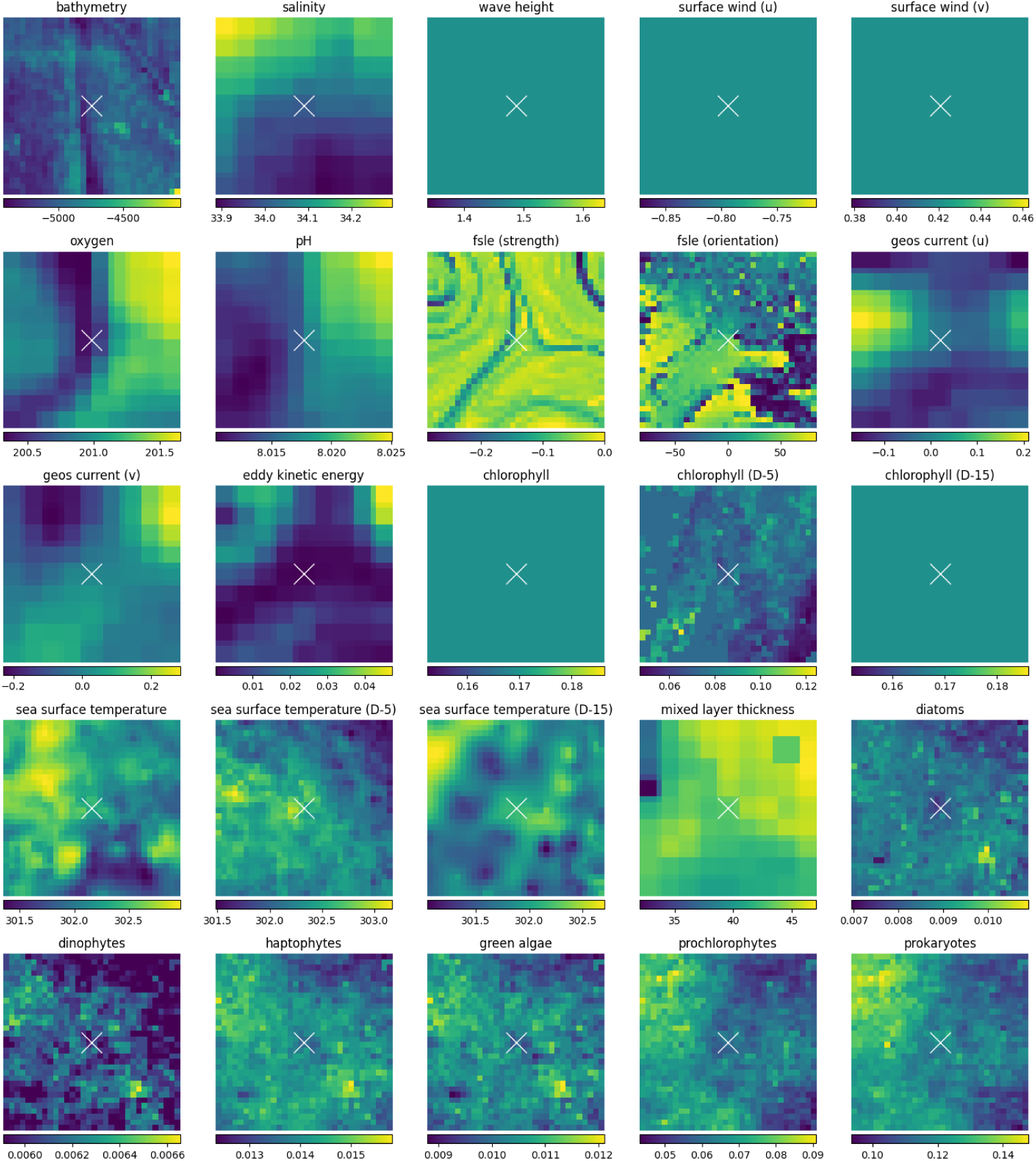
Environmental variables around the point of coordinates –14.389°S, 78.918°E on March 20th, 2021.

### 2.4 Training the model

The modelling technique that we describe in this study was developed for plant species distributions [24]. We used the *Malpolon* framework [43] after some adaptations to our use case. It was built on top of PyTorch [44] and PyTorch Lightning [45] frameworks.

The *Malpolon* framework implements a convolutional network with the *resnet50* feature extractor [22]. We adapted it to use 29 inputs channels and 38 numerical outputs converted to relative probabilities by a Softmax function. Only the first (convolutional) layer and the last (linear + Softmax) layer were altered to adapt the number of inputs and outputs. It was trained from scratch in two sessions: one with a .1 learning rate and another one with a .01 learning rate to fine-tune the weights. We used a Binary Cross Entropy loss (averaged over the taxa and the training batch, weighted by taxa sample sizes).

Treating the problem as a classification task allows estimating the conditional probability of y (the observed species) given that an observation has been made in the environment x. It has the advantage to be (asymptotically) invariant to the spatial sampling effort but it is sensitive to the taxonomic reporting bias (the fact that some species are more observed than others) [46]. In the absence of taxonomic reporting bias, the estimated probabilities would converge to the relative probability of each species given the environment. This is why sample size weights were used in the loss to compensate for this particular bias. Mapping those probabilities is thus equivalent to mapping the species suitability relatively to the other species suitability. It can be related to the “target-group” approach for generating pseudo-absences.

Therefore, the target probabilities for training were set using one-hot encoding, *i.e.* a one for the observed species and zeroes for all others. This follows the principle of assumed negatives [47]: we assume that only the observed species is present at the point of observation. This equates to considering the pseudo-absences of the other species, but it does not prevent co-occurrences. Indeed, if two species are present in the same environmental conditions, training will push the model towards a 50%-50% prediction.

### 2.5 Evaluation metrics and performance assessment

Training data were randomly split into three sets: training (60%), validation (20%) and test (20%). The validation set was used to assess and improve performance during the training phase, while the test set was used after training to compute the final performance of the model, on data that it had never seen before.

In order to assess the advantage of using a convolutional model, we removed geographical input data and retrained the same model from scratch, using only the center value of each patch, for each variable (Punctual-DNN [24]). Since the tiles are 32×32 pixels, we had to use the average of the four center values. We then computed the same accuracy metrics as with our main model; they are displayed side by side in the Results Section.

### 2.6 From predicted probabilities to distribution maps

After training the model, we used it on new data to generate distribution maps. As the environmental data download phase can be quite slow, we had to choose to focus either on time extent or spatial extent, but not both at the same time. This is why we chose to compute two different outputs:

- Global species distribution maps at four dates in 2021.
- Regional species distribution maps for the Southwestern Indian Ocean at 53 dates in 2021.

It is important to note that the model may be used at any date in any area; the only limitation is the availability of environmental data and the time required to download them.

Two grids covering both areas were generated, with an approximate 100 km stride. They comprised 36,506 points for the global oceans and 3,001 for the Southwestern Indian Ocean. Environmental data were downloaded for each of these points and run through the model, which led to 38 predicted probabilities for each of these points, at each requested date. For each taxon, these probabilities were interpolated over the whole area to generate rasterized distribution maps. We used cubic interpolation to generate outputs with a 3600×1800 pixels resolution for the World maps and 800×800 pixels for the Western Indian Ocean maps.

It is worth noting that since we are working with probabilities that are relative to our choice of studied taxa (because of the softmax layer), the absolute values have little purpose. Therefore no scale is provided for all distribution maps: they should be interpreted relatively to one another, across species, time or space.

### 2.7 Influence of variables

To study the influence of variables, a new model was trained after removing chlorophyll and sea surface temperature at D-5 and D-15, as well as Eddy Kinetic Energy. Indeed, these layers are highly correlated with chlorophyll and sea surface temperature on the day of occurrence and geostrophic current respectively.

It is worth noting that this model has almost the same accuracy (69.08%) as the previously described one (69.15%), which shows that the 5 variables that were removed have very little influence on the classification.

Afterwards, the most determining variables were calculated using the integrated gradients method, which is a way to estimate the gradients of the scores with regard to the inputs, therefore gauging the importance of each input data point [48]. They were calculated for all points on the world grid at the four dates of 2021, using the Captum python package [49]. They were then aggregated over the whole study area (sum of absolute values) to represent variable importance over all the world oceans. Finally, to analyze this in a taxon-specific way, this process was repeated for each taxon on a random sample (N=1000) of the points where the taxon was the top prediction.

## 3. Results

### 3.1 Performance of the model

The accuracy of the final version of the model was 69%, which means that in 69% of cases, the most likely taxon according to the model was the same as the one that was actually observed. The corresponding score for the Punctual-DNN is significantly lower: 63%. See Table 3 for more complete accuracy results. These metrics prove the benefit of using spatial data, as hypothesized in the Introduction. Although the difference in scores is quite small, percentage points closer to 100% are much harder to gain than those close to 0%. Indeed, they represent the most difficult predictions.

**Table 3.** Probability that the observed taxon is among the Top N predictions of the model, for 11 values of N.

| Top N | Probability<br>(Spatial input) | Probability<br>(Punctual input) |
| --- | --- | --- |
| 1 | 69.15% | 63.34% |
| 2 | 83.19% | 78.77% |
| 3 | 89.13% | 85.85% |
| 4 | 92.83% | 90.26% |
| 5 | 95.13% | 93.18% |
| 6 | 96.63% | 95.29% |
| 7 | 97.48% | 96.45% |
| 8 | 97.99% | 97.28% |
| 9 | 98.43% | 97.87% |
| 10 | 98.75% | 98.29% |
| 38 | 100.00% | 100.00% |

All these statistics were computed on the test set, *i.e.* on data that was never used before.

A confusion matrix was computed on the test data set and is shown in Figure 4. It shows that some taxa were well predicted by the model (the top two being *Aptenodytes forsteri* and *Mobula alfredi*). Others were harder to predict, the worst two being *Istiompax indica* and *Carcharhinus longimanus*. These two are among the taxa with the fewest occurrences, which could explain this result.

**Figure 4.**
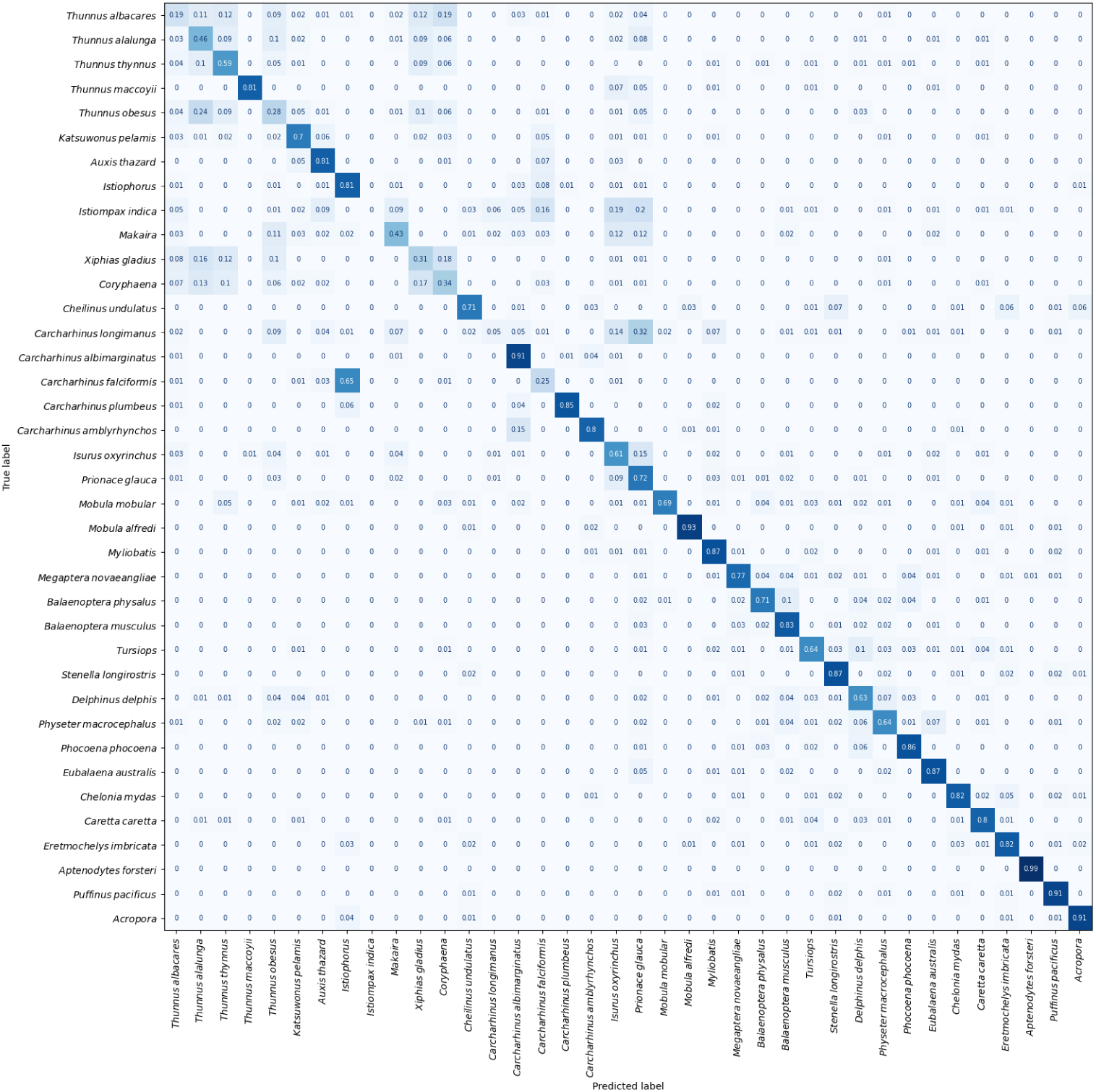
The confusion matrix shows the predictions of the model on the test data set, for each actually observed taxon. Cell darkness is proportional to cross-probability.

### 3.2 Presentation of the species distributions maps

#### 3.2.1 Global oceans

Distribution maps were calculated on four dates, all in 2021, corresponding to both solstices and both equinoxes, for the thirty-eight taxa. These maps represent the probability of presence *among the 38 studied genera*. Figure 5 shows these maps for three species on the spring equinox. All 152 distribution maps are available online [50].

**Figure 5.**
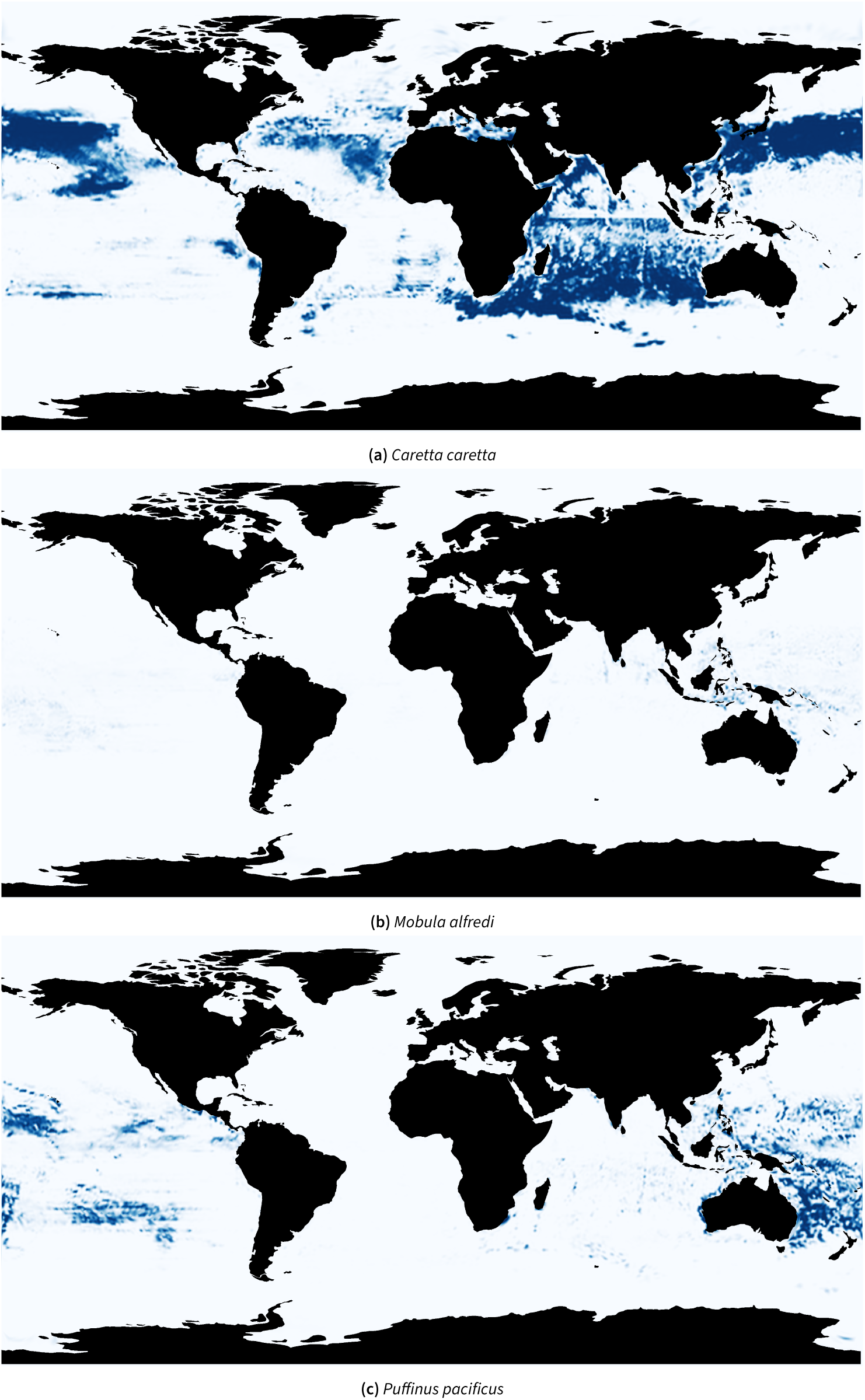
Examples of distribution maps on March 20th, 2021, chosen to further discuss some interesting and contrasting patterns. All maps are available publicly [50].

#### 3.2.2 Southwestern Indian Ocean

In this case, distribution maps were calculated each week of 2021, for the 38 taxa. To make visualization easier, they were exported as animated GIFs that are available online [51]. Again, these maps represent the probability of presence *among the 38 studied genera*. An example for *Prionace glauca* is shown in Figure 6.

**Figure 6.**
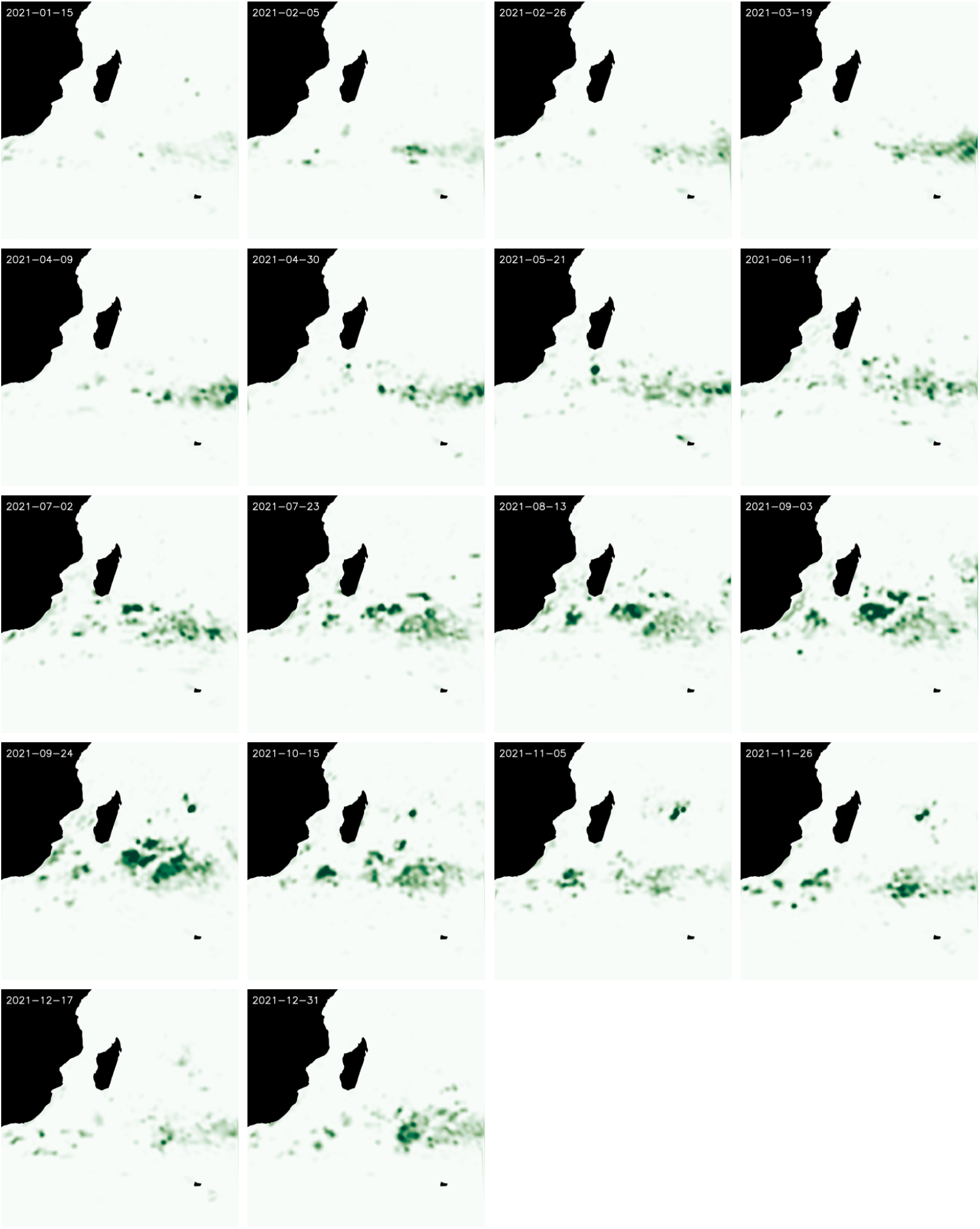
Distribution maps for *Prionace glauca* every three weeks of 2021. The scale is different from other figures to improve visibility, hence the change in colours.

### 3.3 Comparison of predicted distribution maps to established maps

Validation of the distribution maps is challenging because existing distribution maps are usually the results of static studies (except sometimes broad seasonal variations). Yet the maps that we produce are dynamic, *i.e.* highly dependent on time, see Figure 6 for instance. Moreover, our maps show presence probabilities relatively to the set of 38 species, which yields results that are different in nature from classic distribution maps. Finally, observation data are spatially biased so they cannot be used for validation either.

We compared some of our distribution maps to established ones, to check for inaccuracies. See Figure 7 for a few examples.

**Figure 7.**
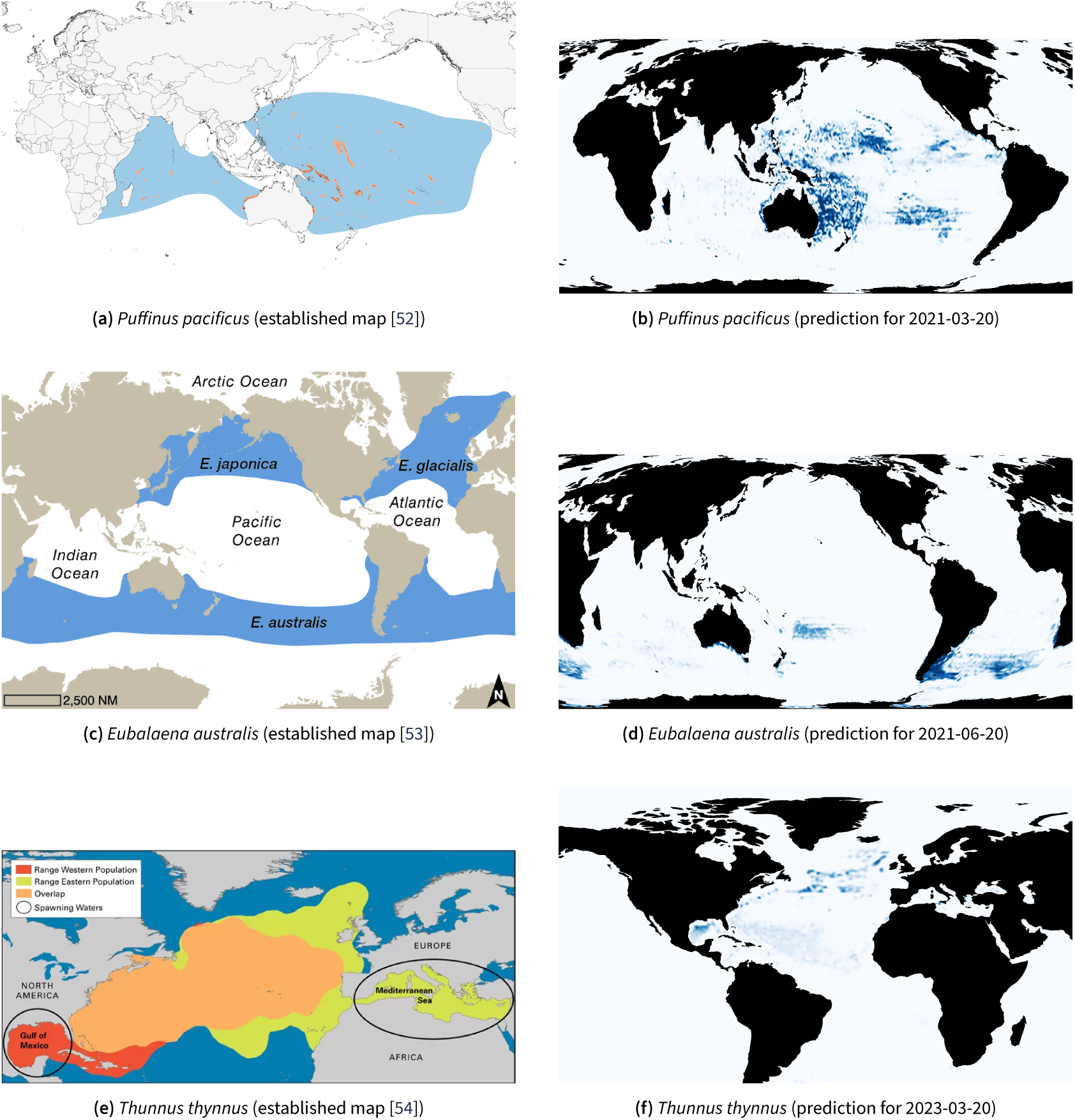
Comparison between established distribution maps (left) [52, 53, 54] and deep-learning generated maps (right). Prediction maps at other dates [50] do not change the interpretation.

#### Puffinus pacificus

The prediction map 7a is consistent with the established one 7b, although it shows a significant difference between the Indian and Pacific oceans. Since established maps are usually binary (presence/absence), the difference we see between the two oceans cannot be (in)validated. It is possible that the Pacific Ocean is more suitable to this species than the Indian Ocean, or the Indian Ocean stock may be an under-represented in our data set. In theory, this should not have impacted our results as the point of the method is to be geography-agnostic. But in reality different stocks may have different reactions to environmental conditions, therefore introducing a correlation with geography (location of the stocks). Perhaps some stocks were under-represented in our training data set.

#### Eubalaena australis

The predicted distribution 7d fits within the known geographical range of *Eubalaena australis* 7c and the map shows a strong disparity of the prediction density within this area. Again, no assumption can be made on the plausibility of the predictions, as this heterogeneity may be caused by temporal variation, or it may not fit reality. As our results are relative probabilities (*i.e.* proportions among all 38 species), variations in one distribution map may also ensue from variations in other species habitat preferences.

#### Thunnus thynnus

The predicted range for *Thunnus thynnus* 7f is within the established range 7e, but it does not include all of it. Specifically, the Mediterranean Sea and the Bay of Biscay are excluded, even though a major population lives in these areas [55]. After checking our input data, this shortcoming can be explained by the under-representation of this population in the occurrences used for training. This will be discussed further in Section 4.2.2.

### 3.4 Analysis of determining variables

Over the predictions for the 2021 Global use case, the most influential variables were finite-size Lyapunov exponents (FSLEs) (strength), sea surface temperature (SST), pH, salinity, FSLEs (orientation) and bathymetry, in this order. See Table 4 for a full accounting of variable influence.

**Table 4.** Statistics of the influence of variables over the 2021 predictions for the whole world (×1, 000, sorted by median). Colour scale 0 to 1. MAD = Median Absolute Deviation.

| Variable | min | 25% | 50% | 75% | max | MAD |
| --- | --- | --- | --- | --- | --- | --- |
| FSLEs (strength) | 0.00 | 0.42 | 0.73 | 1.21 | 6.88 | 0.36 |
| SST | 0.00 | 0.17 | 0.44 | 0.95 | 5.13 | 0.33 |
| pH | 0.00 | 0.19 | 0.38 | 0.72 | 5.04 | 0.23 |
| Salinity | 0.00 | 0.19 | 0.32 | 0.63 | 3.56 | 0.17 |
| FSLEs (orientation) | 0.00 | 0.16 | 0.30 | 0.56 | 3.85 | 0.17 |
| Bathymetry | 0.00 | 0.11 | 0.28 | 0.66 | 8.39 | 0.22 |
| Geos. Current (u) | 0.00 | 0.13 | 0.24 | 0.41 | 3.25 | 0.12 |
| Geos. Current (v) | 0.00 | 0.14 | 0.24 | 0.40 | 2.93 | 0.12 |
| Surface wind (v) | 0.00 | 0.13 | 0.23 | 0.38 | 1.48 | 0.11 |
| Surface wind (u) | 0.00 | 0.12 | 0.20 | 0.34 | 1.43 | 0.10 |
| Oxygen | 0.00 | 0.05 | 0.20 | 0.61 | 4.26 | 0.18 |
| Wave height | 0.00 | 0.07 | 0.11 | 0.19 | 0.83 | 0.06 |
| Mixed layer thickness | 0.00 | 0.03 | 0.09 | 0.28 | 3.83 | 0.07 |
| Pacific Ocean | 0.00 | 0.00 | 0.08 | 0.57 | 3.29 | 0.08 |
| Green algae | 0.00 | 0.01 | 0.03 | 0.06 | 2.88 | 0.02 |
| Prochlorophytes | 0.00 | 0.01 | 0.03 | 0.06 | 3.62 | 0.02 |
| Haptophytes | 0.00 | 0.01 | 0.02 | 0.06 | 2.93 | 0.02 |
| Prokaryotes | 0.00 | 0.01 | 0.01 | 0.03 | 2.24 | 0.01 |
| Chlorophyll | 0.00 | 0.00 | 0.01 | 0.02 | 1.23 | 0.01 |
| Dinophytes | 0.00 | 0.00 | 0.01 | 0.02 | 3.14 | 0.01 |
| Diatoms | 0.00 | 0.00 | 0.00 | 0.02 | 1.96 | 0.00 |
| North hemisphere | 0.00 | 0.00 | 0.00 | 0.48 | 3.84 | 0.00 |
| Atlantic Ocean | 0.00 | 0.00 | 0.00 | 0.12 | 3.27 | 0.00 |
| Indian Ocean | 0.00 | 0.00 | 0.00 | 0.00 | 3.35 | 0.00 |

Figure 8 shows the median integrated gradient for each taxon. While FSLEs and SST are the most important variables overall, this chart reveals the diversity of correlations between taxa and predictors. For example, bathymetry was the most important predictor for 6 taxa. Another interesting observation is that phytoplankton data was used significantly for one taxon only: *Thunnus thynnus*.

**Figure 8.**
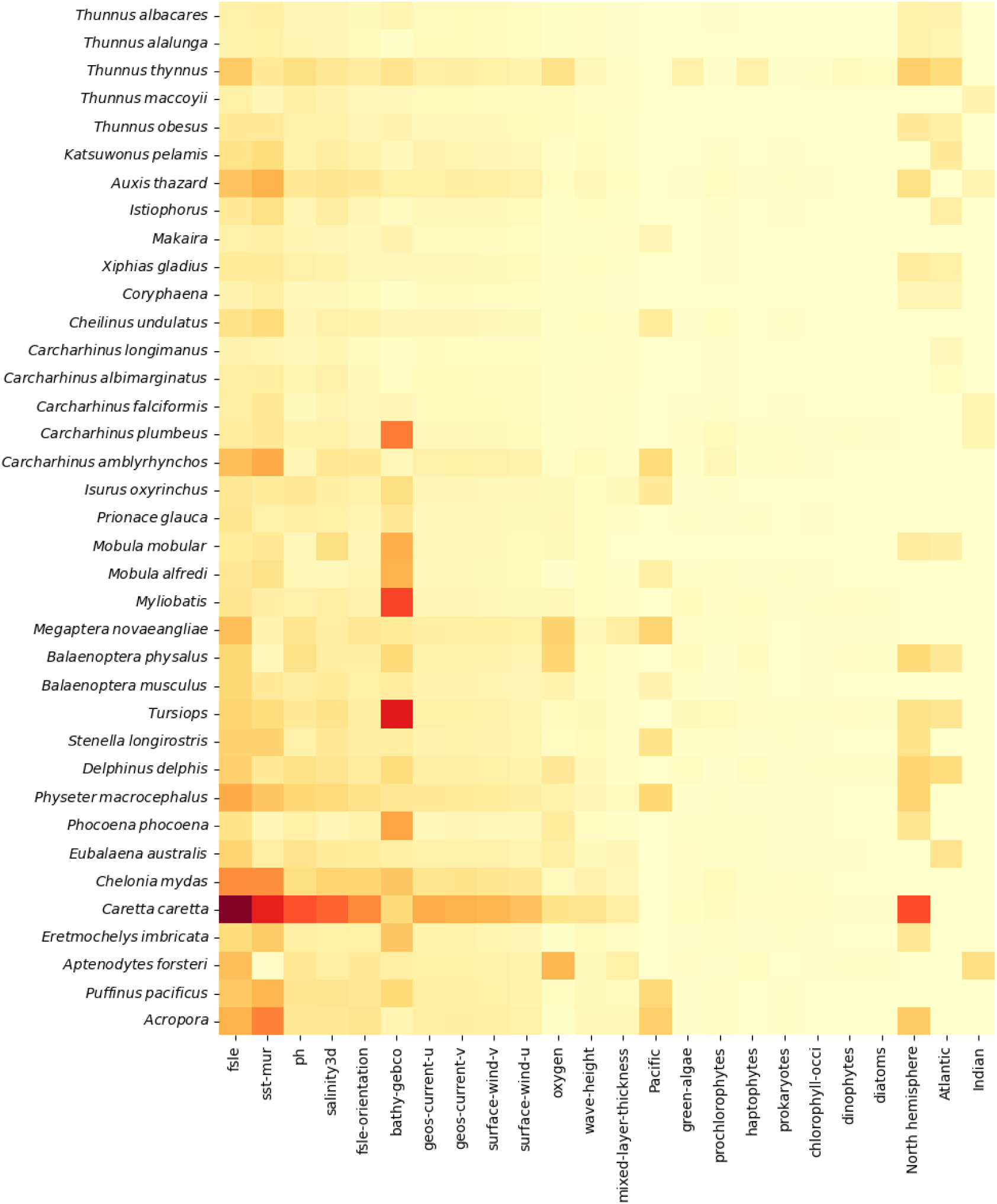
Variables that had the most influence on the determination of each taxon presence (darker = stronger influence). NB: *Istiompax indica* is not present in this chart as it was never predicted to be the most likely taxon.

## 4. Discussion

### 4.1 Ecological interpretation of the results, implications for offshore species distributions

The variables that were identified as important are coherent with past research. Specifically, FSLEs were identified as a particularly important predictor of movement for top marine predators [56]. Sea surface temperature was also expected to be an important predictor, since it has important physiological consequences and is therefore the most frequently used descriptor in marine SDMs [21] and was identified as the most relevant factor in an SDM review [57].

This study also demonstrates a high sensitivity to temporal variations in environmental conditions, as shown in Figure 6. This highlights the need for distribution models of fast-moving species to consider these variations and is coherent with previous findings [19, 21].

We noticed some surprises in influential variables: bathymetry was not a good predictor of *Acropora* coral distribution, which is contradictory with their need for light. A possible explanation is that the model may have used other variables as a proxy for low depths. This could be a legitimate and expected behavior, or overfitting due to auto-correlated training data, which is discussed in the next section.

### 4.2 Benefits and limitations of using deep learning for SDMs in the open ocean

This method holds promise in helping researchers uncover new correlations between the oceanic conditions and species distributions: implicit feature extraction allows the use of more numerous and more complex features. Indeed we showed that the convolutional part of the model was taking advantage of spatial data, which lead to significantly higher accuracy than using data only from the point of occurrence. In this study, we showed the variables that had the most influence on average. This needs to be complemented by a deeper study of the nature of the determining features.

We noted three main limitations of our method, namely performance metrics, biases in the input data and some undetected patterns.

#### 4.2.1 Accuracy metric

The present model is a classifier and as such, it behaves differently from usual SDMs. In particular, outputs are predicted probabilities, relatively to the set of 38 taxa. Therefore, prediction maps cannot be interpreted as the usual results of SDMs. For example, if a species is obviously present in some environmental conditions, probabilities for other species will be lower. A solution to this shortcoming would be to include a large number of species into the study (*e.g.* 4,520 in [24]), which is a priority for any reproduction of this work.

The accuracy of the model could still be improved, depending on the ecological feasibility. Indeed, as individuals, members of a species may explore, behave erratically, or in any other way exercise their free will or at least their individual preference [58]. Their response to environmental predictors may even depend on the environment itself [59]. As such, dynamic SDMs will never provide a perfect prediction of their distribution.

Furthermore, if some species are frequently seen together, the model cannot discriminate between the two. In that case, this uncertainty will show as a .5 mistake rate even though it is the correct result. A way to improve the final accuracy score would be to group species by habitat preferences, but this would remove the possibility of studying differences between such species. For example, the confusion matrix in Figure 4 shows that *Xiphias gladius* and *Coryphaena* are often predicted instead of each other. This may be the result of these two taxa having similar habitats, and the low resulting score does not necessarily mean that the predictions are wrong.

Consequently, accuracy is not an ideal metric for this use case, as we use a classifier to bypass the scarcity of training data, in particular the fact that almost all available data are presence-only. This should not be a deterrent as previous research showed that presence-only occurrence data could yield satisfactory results [60]. This is why we provide a Top-N score in Table 3 for a more complete performance assessment.

#### 4.2.2 Observer bias

Most observation data in the open ocean come from fishing vessels, which target certain species. This causes observations to mostly include target species or frequently associated species. Furthermore, fishing boats tend to target some areas based on outputs of fishing guidance models so it creates an artificial correlation between the parameters used in these models and the presence of animals [61].

The fact that the model has limited access to geographical information (only hemisphere and ocean basin) partly compensates for sampling effort heterogeneity. Indeed, only environmental conditions guide the predictions. But this is not flawless when various stocks of the same species have different behavior relatively to environmental conditions. This is the case of *Thunnus thynnus* which has two separate stocks (West and East Atlantic) [62]. When sampling is biased between stocks, the model might not fully learn the various responses to environmental responses. This explains why the model failed to extrapolate from West Atlantic data and to predict high probabilities in the Mediterranean Sea and the Bay of Biscay.

Finally, some data may come from scientific tracking of individual animals, so these individuals may be over-represented in our data and reflect their preferences rather than the general tendency of their species. The large amount of occurrences that we used help tackle this bias.

These biases would be better tackled with more available data, which is a serious issue in the open ocean. Little data is produced relative to the size of the oceans and a large part of this data is not shared publicly. More data is key to better models and more trustworthy distribution maps, as previous research even showed that more data was more important than spatial bias in our context [63].

#### 4.2.3 Undetected patterns

Detection of seasonal migrations is incomplete. For instance, we should see the *Megaptera novaeangliae* distribution spreading north during the southern winter [64]. The model also did not catch the *Thunnus thynnus* seasonal spawning in summer in the Mediterranean Sea.

The causes of these shortcomings are unclear, so we offer a range of possible explanations and ways to improve the present method, in the hope that these will help future research obtain higher quality results.

### 4.3 Suggestions to further improve the modelling methods

#### 4.3.1 Occurrence data

As a first experiment, occurrence data were selected randomly for this study. Even though the aim should not be to have a perfect fit between observation data and model predictions, observer bias could be reduced by selecting data sources more appropriately. In particular, redundant data sets should be avoided and more importance should be given to the diversity of sampling methods.

As previously discussed, this method would probably benefit from including a large number of taxa. In particular, planktonic species may prove valuable as they are less prone to sampling biases [65] and data sets are widely available [66].

Further, It could be interesting to run the model on two separate occurrences data sets: one with fast moving species only and the other with sedentary or sessile species only. This would allow testing the efficiency of dynamic SDMs in two different contexts: real-time environmental conditions preferences and long-term distribution shifts, respectively.

#### 4.3.2 Environmental data

For some variables, it could be beneficial to use other sources. For example the chlorophyll data that we used was quite incomplete and using the Copernicus product instead [42] could yield better results. It has also been suggested to include 3D environmental data, as most variables vary with depth and occurrence data are not limited to the surface [67]. Such data could easily be included in the input data with no change to our method. Finally, additional data may be beneficial, in particular the distance to the nearest coastline or level of anthropisation.

The encoding of variables could also be experimented with. In particular, vectors (FSLEs, wind and current) could be encoded with three variables instead of two: strength, cos(angle) and sin(angle). This would make the North-South and East-West components, as well as the total strength explicit.

#### 4.3.3 Model training

Several choices could be made differently during the training phase. For example, *Acropora* presence is predicted in the open waters of the Pacific (too much to correspond to Pacific atolls), even though it is only present at very low depths in both the training and testing data sets. This may be the result of overfitting, due to autocorrelation between the training and validation data sets [68]. This is not visible in the covariance matrix because in that case the testing data set is also correlated with the two others. To remedy this, the split between the training, validation and testing data sets could be more sophisticated, by using block cross-validation or withholding a region/period only for testing, or even more complex methods such as adding a time lag to some observations [69].

Second, although we experimented with loss functions, this can be continued to try and find alternatives more adapted to this context.

#### 4.3.4 Removing artefacts

Three types of graphical artefacts are present in our results:

- The sharp divide between geographical areas caused by our binary geographical variables (see the Indian Ocean in Figure 5a for instance). Ideally, for a given species, only barriers outside of its range should be significant and therefore barriers should not affect the maps. Indeed, we included these artificial barriers as proxies of the historical zoogeographical barriers to colonization [70]. But the imperfections of the model and the low number of species (see third point) contributed to this flaw in the maps. This type of artefact could be mitigated by using a gradient between corresponding binary variables in the zones where theses areas meet. A more drastic solution would be to fully remove the binary geographical variables. This would imply either **1.** using the model only on smaller regions with full connectivity or **2.** accepting that the model predicts theoretical habitat suitability, independently from actual species presence.
- The spotted aspect of the map, which shows that the model uses small scale features for its prediction. Further investigations need to be conducted to determine whether they are justified by the environmental preferences of taxa or if they are the result of overfitting.
- The third one is less visible, but it is a consequence of our predicted probabilities being relative to the 38 species. When an area is favorable to species A, but species B dominates in part of this area, the distribution map for species A shows variations over the area (regardless of actual variations in suitability to species A). Figure 9 shows an example of this phenomenon where the presence of *Katsuwonus pelamis* causes a hole in the distribution of *Caretta caretta*. This makes it harder to interpret the maps, and a solution would be to increase the number of species, as previously suggested in other sections. This way the variation of suitability to one species would only marginally influence probabilities for other species. The present study shows that *N* = 38 is not enough to avoid this type of artefact.

**Figure 9.**
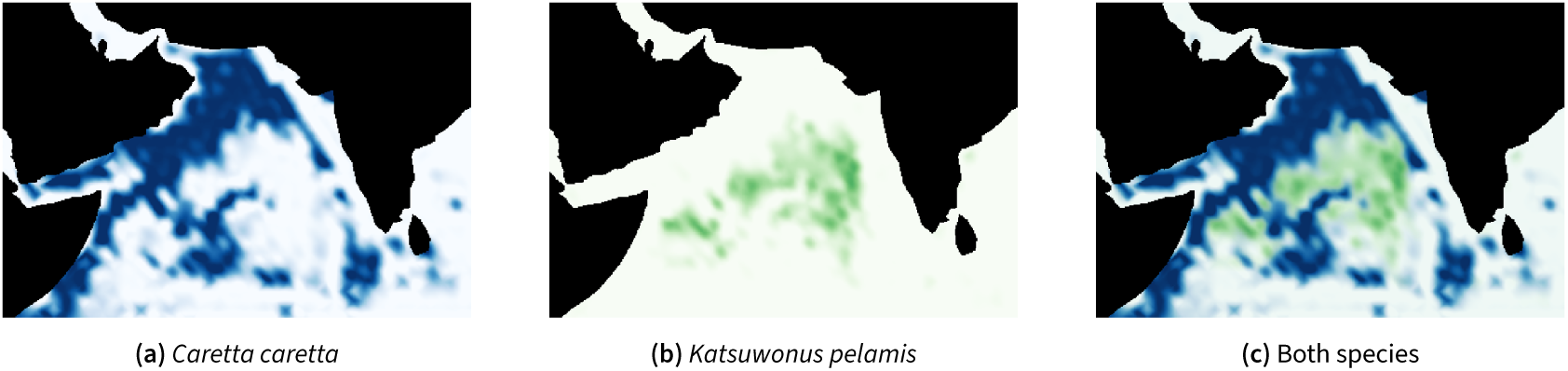
Example of how one species distribution can influence results for an other.

#### 4.3.5 Other use cases

The present method could be used at different scales, in particular in coastal areas. This would require a significant change in the input variables, as the resolution of globally available environmental data is a limiting factor. They could be replaced by satellite or drone images, as well as locally available (more precise) environmental data.

## 5. Conclusion

### 5.1 Main findings and their significance

Dynamic SDMs provide a way to estimate species presence at all dates and all areas, provided environmental data is available. In addition, the present method leverages environmental data around occurrences, including complex patterns. While the dynamic nature makes it difficult to judge accuracy (available reference data are static), it provides a baseline that can be calculated for any species (that have enough existing observations). Researchers working on terrestrial plants have also shown that such models may be used to infer species distribution for rare species, by extrapolating results from co-occurring species [24].

### 5.2 Implications for management and conservation of offshore species

We hope this method will be developed further and used on other endangered species, together with existing methods and field observation. The technique that we presented would be especially useful in the hands of scientists who are experts in the life cycle of specific species. It would help them increase scientific knowledge of their distributions, which is essential for decision-makers to target areas of interest for conservation. Our method would also greatly benefit from their input, as we do not have the species-specific expertise that is necessary to fine-tune training and predictions.

### 5.3 Recommendations for future research and potential applications

While the accuracy of our distribution maps is difficult to assess, there is exceptional room for improvement and further research. All the blocks in Figure 1 can be modified, either to adapt the process to a different use case or to try to improve the quality of the results. Here are some examples of potential changes:

- To study other species, the initial choice of species can be changed, for example, to focus on sedentary species or a specific area.
- To improve accuracy, the occurrence data may be selected in other ways that are not random.
- To investigate the influence of other variables, they may be added to the variable set.
- To study the long-term effect of environmental conditions, some variables may be included with a longer time lag such as months or years.

Finally, we expect that experts of different taxa will rightfully criticize the maps we provide. We wish they did not have to, but developing such a model inevitably includes a trial and error phase, so we welcome their remarks which will lead to investigating issues and proposing improvements for subsequent studies.

The results we presented in this article are a small part of what can be achieved with this model. Many other scientific questions can be investigated both with the model we provide (already trained) or with other models trained with the same method.

## Acknowledgement

This study would not have been possible without Sylvain Poulain and his support in setting up docker images and GPU calculation environments.

We thank Hervé Demarcq for sharing his expertise on environmental data products and Emmanuel Chassot for his invaluable insights into pelagic fishes and species distribution models.

This study was significantly improved thanks to numerous and thorough reviews from Jean-Olivier Irisson, Sakina-Dorothee Ayata and a third reviewer who chose to remain anonymous. We are also grateful for the platform provided by PCI Ecology for ethical reviewing and publishing, and the comments by recommender François Munoz.

## Funding statement

This study was conducted as part of the G2OI project, co-financed by the European Union, the Reunion region and the French Republic.

## Open data statement

### Code

The code that was used to prepare the data, train the model and export the outputs is available on GitHub in the IRDG2OI/deep-sdm-oceans repository and on Zenodo [71].

### Input data

The input data include the CSV file describing the geographical points, the standardized numpy arrays of corresponding environmental data and the standardization factors. They are available on Zenodo [72] for each use case:

- Training data (includes train+validation+test)
- Prediction data for the world at 4 dates
- Prediction data for the Western Indian Ocean at 53 dates

### Modelling

We provide the model checkpoint and configuration file [73], so researchers can make predictions with the presently described model.

We also provide the code that was used for training so researchers can adapt it to their needs and retrain a new model [71]. It consists of Python files based on a custom version of Malpolon.

### Results

The distribution maps were uploaded to Zenodo for easy visualisation, in two repositories:

- Global predictions as PNGs and GeoTIFFs [50]
- Western Indian Ocean predictions as GIFs and GeoTIFFs [51]

## Notes

https://github.com/IRDG2OI/deep-sdm-oceans

https://doi.org/10.5281/zenodo.8188512

https://doi.org/10.5281/zenodo.8202914

https://doi.org/10.5281/zenodo.8202261

https://doi.org/10.5281/zenodo.8202055

### Competing Interest Statement

The authors have declared no competing interest.

### Summary of Updates

This version of the manuscript has been revised to take into account the follow-up comments made by the three PCI Ecology reviewers. In particular, we expanded on the discussion. A detailed change log will be provided on the PCI Ecology website.

